# Oligomeric hypercomplex of the complete oxidative phosphorylation system in heart mitochondria

**DOI:** 10.1101/2020.08.17.253922

**Authors:** S.V. Nesterov, Yu.M. Chesnokov, R.A. Kamyshinsky, A.A Panteleeva, K.G. Lyamzaev, R.G. Vasilov, L.S. Yaguzhinsky

**Affiliations:** Moscow Institute of Physics and Techonology, 141701 Dolgoprudny, Moscow Region, Russia; Institute of Cytochemistry and Molecular Pharmacology, 115404 Moscow, Russia; National Research Center « Kurchatov Institute», 123182 Moscow, Russia; Federal Scientific Research Centre “Crystallography and Photonics” of Russian Academy of Sciences, 119333 Moscow, Russia; Belozersky Research Institute for Physico-Chemical Biology, Lomonosov Moscow State University, 119992 Moscow, Russia

**Keywords:** Mitochondria, supercomplex, respirasome, ATP synthase, oxidative phosphorylation, cryo-EM

## Abstract

The existence of a complete oxidative phosphorylation system supercomplex including both electron transport system and ATP synthases has long been assumed based on functional evidence. However, no conclusive structural confirmation has been obtained. In this study cryo-electron tomography was used to reveal the supramolecular architecture of the rat heart mitochondria cristae. We show that rows of respiratory chain supercomplexes are connected with rows of ATP synthases forming the oligomeric hypercomplex. The discovered hypercomplexes may increase the effectiveness of oxidative phosphorylation ensuring a direct proton transfer from pumps to ATP synthase along the membrane without energy dissipation.

## Introduction

Respiratory supercomplexes (respirasomes) in mitochondria are the keystones ensuring the efficient electron transport and the absence of energy leakage. At the same time, the energy transfer to the final stage of oxidative phosphorylation system (OXPHOS) requires protons rather than electrons. However, while the coupling between proton pumps and ATP synthase has been proposed based on functional analysis [1] and *in vivo* pH measurements [2], the existence of complete OXPHOS supercomplexes – hypercomplexes – was not confirmed. Here, we aim to provide the first structural evidence for the presence of hypercomplexes in mitochondria.

After transmembrane transfer protons do not immediately detach from the membrane [3] due to the existence of kinetic barrier [4,5], therefore OXPHOS clustering can be energetically beneficial. Protons on the membrane surface have a high lateral mobility [6–9], which enables their transfer for the short distances along the membrane without detachment into the bulk phase. The induction of ATP synthesis by the excess protons on the interface boundary was shown for the first time in the octane-water model system [10]. Later the participation of laterally transferred protons on ATP synthesis was reported in mitochondria and mitoplasts [11,12]. Keeping protons on the membrane surface allows mitochondria to synthetize ATP under lower pH gradient [2]. Recently, it was confirmed that the distance between the proton pump and ATP synthase affects the speed and efficiency of ATP synthesis [13]. These data support the idea that for reaching the maximal efficiency all parts of the OXPHOS should be clustered.

The existence of respiratory supercomplexes was shown for the first time in bacteria [14] and later they were detected in different species by inhibitory analysis [15] and by blue-native gel electrophoresis with mild detergents [16,17]. The last method has also showed that the long-established functional link between succinate dehydrogenase and ATP synthase [1] is accompanied by a structural link [18]. It has also confirmed the docking of some mitochondrial dehydrogenases with complex I [19], which was previously demonstrated in the isolated enzymes [20,21]. By now, it has become generally accepted that at least part of the OXPHOS can be organized as a supercomplex [22,23]. All previously described OXPHOS supercomplexes can be divided into two groups. The first is called respirasome and includes respiratory chain complexes in different proportions [24–27]. The second is called ATP-synthasome and includes ATP synthase, phosphate and ADP transporters [28–30]. The direct structural coupling of these two subsystems (producer and consumer of excess protons) has not been revealed yet.

To solve this long standing problem, we used cryo-electron tomography (cryo-ET). The resolution of single particle cryo-ET (SPT) is limited by the thickness of the sample, therefore, we studied the samples obtained through self-destruction of the rat heart mitochondria isolated by the standard protocol. The mitochondria were not exposed to any influences that could have seriously violated their native lipid and protein organization, such as osmotic shock, detergents or centrifugation prior to flash-freezing. The implemented approach allowed to determine different mitochondrial complexes localized in the minimally damaged inner mitochondrial membranes, as well as to obtain their structures with a satisfactory resolution.

## Methods

### Mitochondria isolation

Adult Wistar rats weighting 180-200 grams (9-10 weeks old) were used to isolate mitochondria. All the experiments using laboratory animals were carried out in accordance with the recommendations of the local ethics committee (in accordance with directive 2010/63/EU). All the reagents used have a high purity and were made by reliable manufacturers (Merck, Sigma-Aldrich, Amresco, Serva). The isolation of mitochondria was carried out according to a standard method with minor modifications. Briefly, the heart ventricles of two rats were sheared with scissors, washed from the blood and homogenized by teflon pestle homogenizer in an isolation medium (220mM mannitol, 50mM sucrose, 5mM EGTA, 0.1% BSA, 30mM HEPES/KOH, pH 7.4). The resulting homogenate was centrifuged at 4°C at 700g for 8 minutes, and then the supernatant was collected and centrifuged at 9000g for 10 minutes. The mitochondrial fraction obtained in the sediment was resuspended in the washing medium (as isolation medium, but without BSA and EGTA) and centrifuged at 5000g. The resulting suspension of heavy mitochondrial fraction was used for cryo-EM sample preparation and blue-native gel electrophoresis.

### Blue native gel electrophoresis

The composition of supercomplexes after the addition of a mild detergent was analyzed by blue native polyacrylamide gel electrophoresis (BN-PAGE), as described by Schägger and von Jagow [31] but with minor modifications. Mitochondrial proteins from the heart tissue were solubilized with recently recrystallized digitonin (1 g/g), then Coomassie blue G-250 (2.5 µL of a 5% stock in 500 mM 6-aminocaproic acid) was added to the samples. BN-PAGE was carried out in gradient 4-15% Bis-Tris gels. Both stacking and resolving gels contained 0.025% digitonin. The gels were run at 4-7°C in the Bio-Rad Mini-PROTEAN Tetra system at 100 - 250 V within 3-6 hours. We used the anode buffer (50 mM Bis-Tris with pH 7.0) and the cathode buffer (50 mM Tricine, 15 mM Bis-Tris and 0.002% Coomassie blue G-250 with pH 7.0). As opposed to the original method for BN-PAGE, we did not change the cathode buffer during electrophoresis and it contained a lower concentration of Coomassie blue. After electrophoresis, the gels were stained with Coomassie blue G-250 (0.05% Сoomassie, 40% ethanol, 10% acetic acid), which was followed by a destaining step. We determined the size of the complexes using the SERVA Native Marker. The presence of complex I in all bands that corresponded to supercomplexes was confirmed by in-gel complex I activity assay [32].

### Sample preparation for cryo-EM

The dense mitochondrial suspension was diluted to a concentration of ~0.3mg/ml in a respiration medium (KCl 80mM, MgCl_2_ 1mM, HEPES/KOH 20mM, KH_2_PO_4_ 10mM, pH 7.4) and stored for 1 hour without exogenous substrates in a closed microtube on ice (about 4°C). Then the mitochondria were slowly heated to 20°C. 10 minutes prior to vitrification, phosphorylation was started by adding 10mM glutamate, 4mM malate, 2mM ADP. Then 20 μl of the obtained sample were mixed with 1 μl gold nanoparticles solution (10 nm Colloidal Gold Labeled Protein A, UMC Utrecht, Netherlands) and 3 μl of the mixture were applied to a glow-discharged (30 s, 25 mA) Lacey Carbon EM grid. After blotting for 2.5 s at 4°C, the grid with the specimen was plunge-frozen into liquid ethane in Vitrobot Mark IV (Thermo Fisher Scientific, USA).

### Cryo-Electron Tomography

The study was carried out with Titan Krios 60-300 TEM/STEM (Thermo Fisher Scientific, USA) cryo-EM, equipped with Falcon II direct electron detector (Thermo Fisher Scientific, USA) and Cs image corrector (CEOS, Germany) at an accelerating voltage of 300 kV. 11 tilt-series of the sample were collected automatically with Tomography software (Thermo Fisher Scientific, USA) in low-dose mode with 18000x magnification (pixel size 3.7 Å) and the defocus value in the range between -6 and -8 μm using bidirectional tilt scheme (0°, -2°, …,-58°,-60°, 2°, 4°, …, 58°, 60°). The accumulated total dose was ~ 100ē/Å^2^.

### Tomogram reconstruction

Cross-correlation alignment and tomographic reconstruction were performed using the IMOD software package [33] by simultaneous iterative reconstruction technique (SIRT) and weighted back-projection (WBP) method [34]. Gold nanoparticles were used as fiducial markers for the alignment of tilt-series projection images. 2 tilt series with the alignment error not exceeding 1 pixel were selected for further processing. The reconstructions obtained by SIRT were used for visual inspection and particle picking, while the WBP reconstructions were used for sub-tomogram averaging.

### Segmentation and sub-tomogram averaging

The coordinates of 100 supercomplexes (“particles”) with clearly visible I_1_ and III_2_ were manually picked from tomographic sections with IMOD and utilized for a *de novo* reconstruction in Relion2 [35] using sub-tomogram averaging (protocol described in [36]), which resulted in 44 Å resolution, as determined by Fourier Shell Correlation (FSC) 0.143 criteria. The same particles were used as positive examples for automated segmentation using convolutional neural network utility [37] in an open-source EMAN2.23 package [38] on 2 times binned data (pixel size 7.4 Å) restored with SIRT. ATP synthases, pyruvate dehydrogenase complexes, gold fiducials, edges of the carbon support and contamination on the grid surface were used as negative examples for neural network training.

To find the coordinates of supercomplexes on segmented tomograms, the “Find particles from segmentation” utility in EMAN2.23 was used. 5821 automatically picked sub-tomograms were extracted from 2 times binned dataset (pixel size 7.4 Å) restored with WBP, aligned to a *de novo* model using 3D auto-refinement (Relion2) and sorted into 3D classes without alignment (τ2_fudge=8). One class with 1706 particles was chosen for further processing. Refinement with low-pass filtered to a 60 Å *de novo* model as reference resulted in 24.6 Å resolution (FSC=0.143) supercomplex density.

Since the densities of IV complexes varied, an additional focused 3D classification (τ2_fudge=20) was performed using mask on complex IV. 944 particles (55%) consisting of I_1_III_2_IV_2_, resulted in 27.7 Å density map (EMD-11605), and 762 particles (45%) corresponding to I_1_III_2_IV_1_, resulted in 29.6 Å density map (EMD-11604).

14.8 Å density of ATP synthase dimers was obtained from the same dataset as described previously [39].

Graphics, final visualization and fitting were performed with UCSF Chimera [40]. The PDB-5xtd, the PDB-1bgy and the PDB-5z62 structures from Protein Data Bank were used for the fitting of I, III and IV complexes, respectively. Mitochondrial membranes were manually segmented with Avizo (Thermo Fisher Scientific, USA) software package.

## Results

### Respirasome structure

After the vitrification procedure, the thinnest areas of the prepared EM grids with fragments of phosphorylating mitochondria, consisting mostly of destroyed mitoplasts and tubular cristae forming a continuous network (Fig. S1), were selected for SPT. On the first step of the research the protein composition of respirasomes and mutual arrangement of its components (complexes I, III, IV) were studied with SPT (Fig. 1). The density maps with 27-30 Å spatial resolution were obtained and two types of respirasomes with different number of complexes IV (Fig. S2) were revealed. Comparison of the obtained results with blue-native electrophoresis (Fig. S3) and pertinent literature [24] indicates that the structure of the observed supercomplexes is well preserved. This is confirmed by a much higher relative content of respirasomes containing two complexes IV in our cryo-ET experiments (more than 50%).

**Fig. 1.**
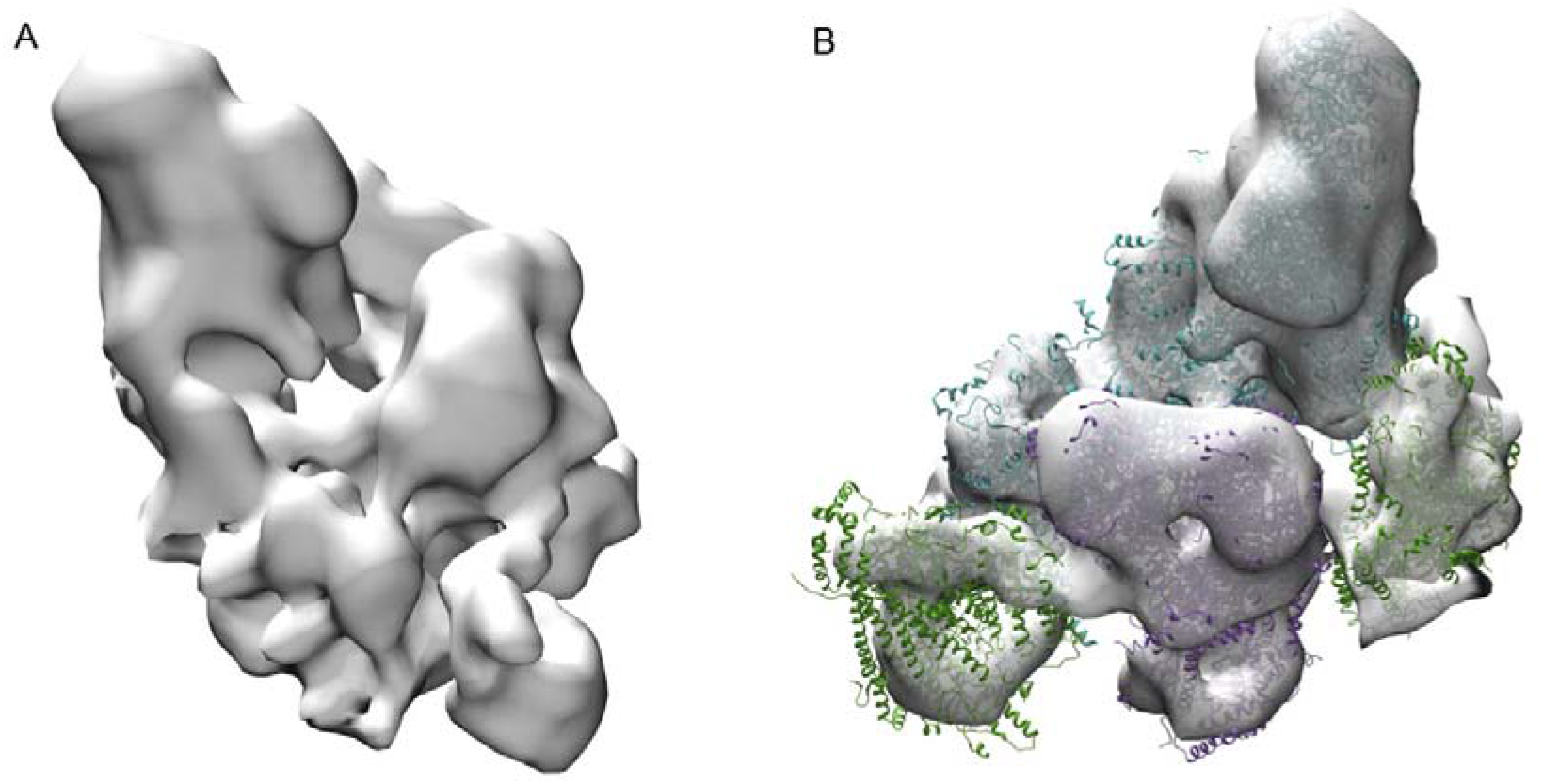
A. Cryo-EM map of the most common type of respirasome (mitochondrial respiratory chain supercomplex). B. 90° rotated cryo-EM map with fitted high-resolution structures (PDB 5z62, 5xtd, 1bjy). Blue - complex I, purple - complex III dimer, green - complex IV.

### The structure of oligomeric OXPHOS hypercomplex

The second step included studying the supramolecular organization of mitochondrial cristae. It has been shown that under experimental conditions the respirasomes and ATP synthases form highly ordered oligomeric elongated structures – the hypercomplexes (Fig. 2A). The cryo-ET data demonstrates that linear structures consisting of respirasomes I_1_III_2_IV_2_ and I_1_III_2_IV_1_ are colocalized with parallel linear structures consisting of ATP synthase oligomers (Fig. 2, 3, S4). The distance between respirasomes and ATP synthases in most of the observed structures is 1-5 nm. Moreover, several areas demonstrate tight docking of ATP synthases with complexes I or IV (comparable with the distance between complexes in respirasomes), which allows considering these areas as the complete OXPHOS supercomplexes. It is essential to point out that in hypercomplexes the elongated transmembrane parts of complexes I are oriented approximately along the row of ATP synthases. While their relative position is undoubtedly not strictly fixed, statistics show that complexes I have a preferred orientation – their elongated membrane parts are virtually never oriented perpendicular to a row of ATPases (Fig. S5). More than that, the row of respirasomes repeats the bend of the ATP synthase row, indicating a possible interaction that holds these structures together. The сryo-ET sections and the 3D reconstructions of cristae are shown in the video (movie 1). It should be noted that in all reconstructions only the reliably identified respirasomes and ATP synthases were demonstrated.

**Fig. 2.**
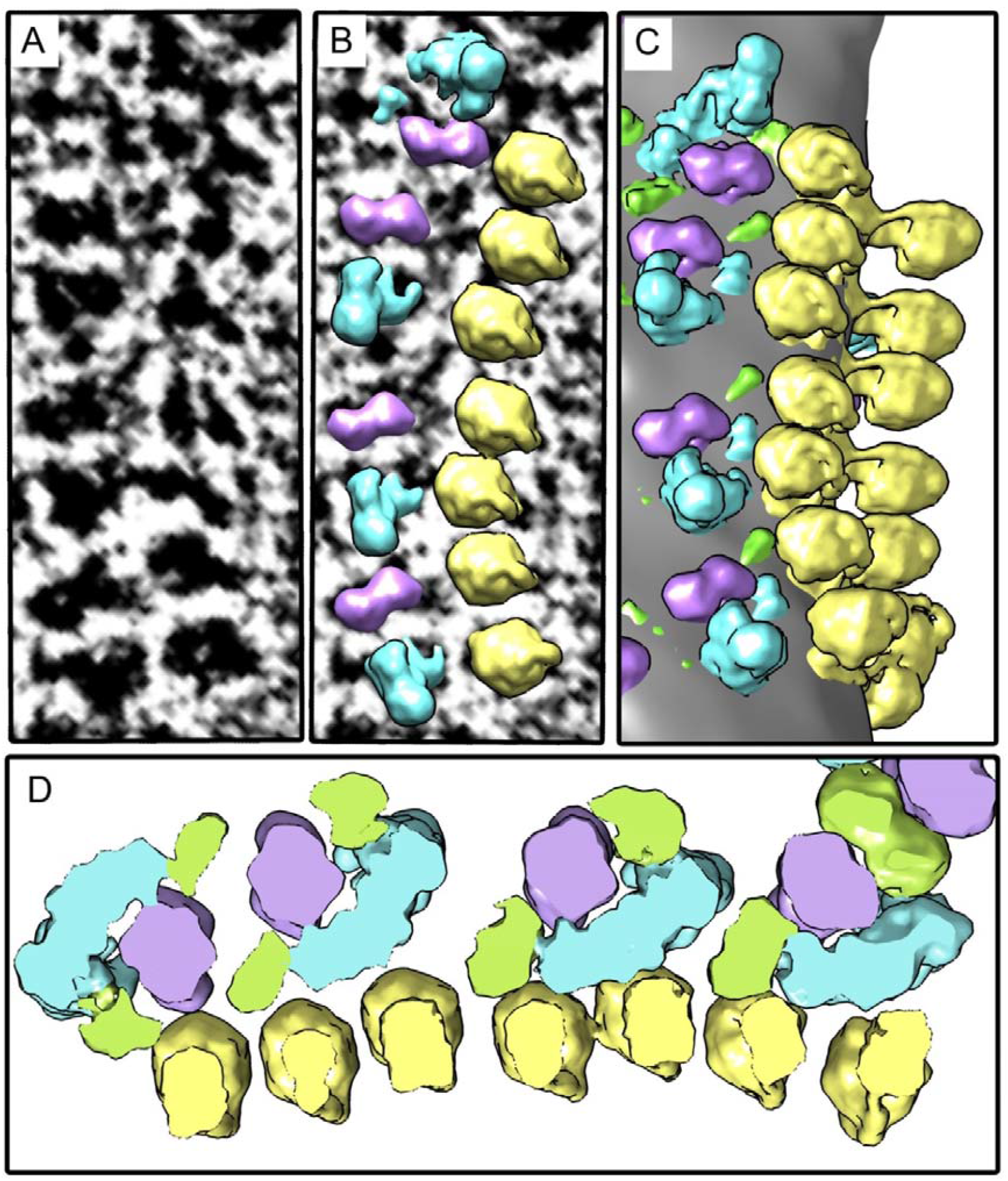
An example of oligomeric linear hypercomplex consisting of tightly docked ATP synthases and respirasomes. Elongated transmembrane parts of complexes I are oriented parallel to the row of ATP synthases. A. Tomographic slice. B. Density maps of complexes I, III_2_ and ATP synthases placed back to the tomogram. C. Surface rendering of the selected area. D. Slice of the density map, view from the intermembrane space. Image D is a bit enlarged in comparison with Fig. A-C. Colors: yellow - ATP synthase, blue - complex I, purple - complex III dimer, green - complex IV.

**Fig. 3.**
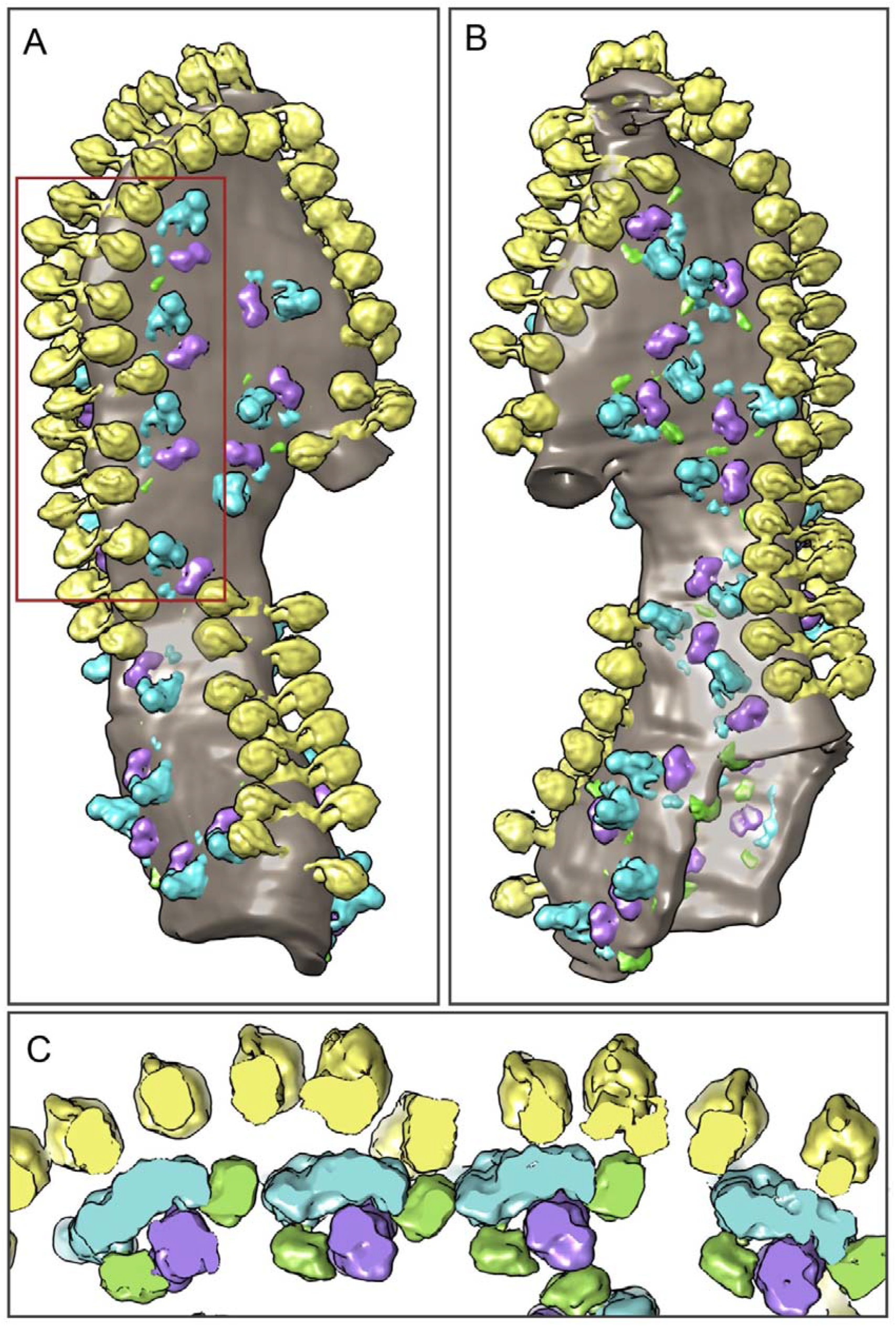
Surface rendering of the mitochondrial crista containing the oligomeric linear hypercomplex of the respiratory chain and ATP synthases. A and B. Views of the crista from the opposite sides (180° rotation around the vertical axis), demonstrating a highly ordered structure of the hypercomplex. C. 3D visualization of respirasomes and the ATP synthases contact zones in the membrane. Slice of the fragment highlighted by the red frame in Fig. A. View from the opposite side of the membrane. Colors: yellow - ATP synthase, blue - complex I, purple - complex III dimer, green - complex IV.

The minimal chain of the OXPHOS supercomplex can be conditionally distinguished from the linear hypercomplexes embracing respirasomes and ATP synthases. It consists of two ATP synthase dimers located on the membrane fold, which are symmetrically docked with the respirasomes at both sides (Fig. S6). It should be emphasized that the contact between enzymes is labile and most likely is caused by the attraction of proteins to the membrane zones with a certain folding or lipid composition. This distinguishes the observed structures from classical supercomplexes with fixed protein contacts.

## Discussion

Cumulatively, our data offer the first structural evidence on the complete OXPHOS supercomplex. Unexpectedly, the coupling of proton pumps and ATP synthases occurs not at the level of individual enzymes, but via attraction of large ordered oligomeric structures that results in the formation of hypercomplexes. The existence of these tightly docked structures still raises the question of their functional significance and operation mechanisms to be discussed further. The distinctive peculiarity of the observed structure is that it allows a direct proton transfer from proton pumps to ATP synthase.

The existence of stable ATP synthase rows in mitochondrial membranes was previously shown by various methods [39,41–43]. The possibility of linear structures formation from respirasomes was assumed earlier on the basis of structural analysis [44], but it was not proved experimentally. Moreover, there was no experimental evidence of a structural connection between respirasomes and ATP synthases.

In the discovered oligomeric hypercomplexes, respirasomes are tightly docked to ATP synthases, which indicates OXPHOS clustering. It remains unclear to what extent the discovered ordered orientation of respirasomes is important for the OXPHOS functioning. It can be assumed that the formation of hypercomplexes occurs by a mechanism similar to the formation of membrane rafts – due to the special curvature and lipid composition of the membrane in the vicinity of ATP synthases. Nevertheless, it appears that the effectiveness of OXPHOS may be controlled by the transition between a highly ordered oligomeric state with minimal proton leaks and a less ordered diffusely distributed state. However, further research is needed to unveil the mechanisms of this regulation and its physiological significance.

## Conclusion

Here a new type of the OXPHOS structural organization was revealed - an oligomeric hypercomplex formed of parallel rows of respirasomes and ATP synthases. The short distance between proton pumps and ATP synthases ensures a direct transfer of hydrogen ions to ATP synthases with maximum speed and minimal leakages. Thus, the discovered hypercomplexes should provide both quickness and efficiency of ATP synthesis which are essential for heart mitochondria under high load conditions. Formation and dissipation of such hypercomplexes can be one of the mechanisms for synchronization of the OXPHOS components and their activity regulation.

## Acknowledgments

This work has been carried out using computing resources of the federal collective usage center Complex for Simulation and Data Processing for Megascience Facilities at NRC “Kurchatov Institute”.

## Author contributions

L.S.Y., R.G.V. designed the study; S.V.N. and L.S.Y. wrote the manuscript, which was commented on by all authors; S.V.N. performed mitochondria preparation and manual annotation of the tomograms; A.A.P., K.G.L. performed blue-native gel electrophoresis; Y.M.C., R.A.K. performed cryo-EM experiments and computer data processing;

## Data and materials availability

All data are available in the manuscript or the supplementary material, electron microscopy density maps are deposited in the EMDB (EMD-11605, EMD-11604).

## Supplementary Materials

Materials and Methods

Figures S1-S6

Movie S1

