## Supplementary figures 1-6 for "Oligomeric hypercomplex of the complete oxidative phosphorylation system in heart mitochondria"

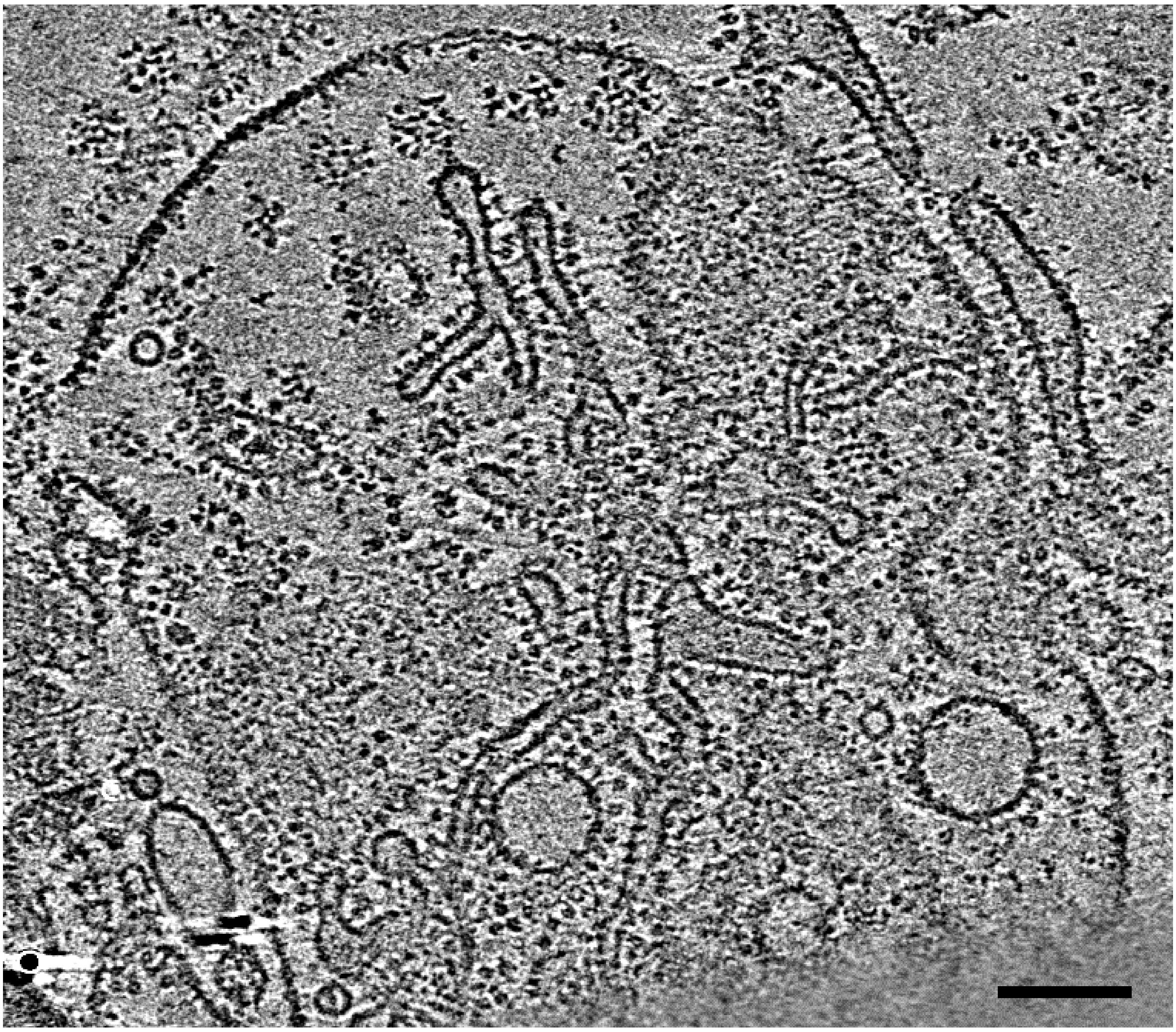

*Figure S1. Tomographic slice of partially destructed heart mitoplast, demonstrating a continuous network formed by tubular cristae. Scale bar is 100 nm.*

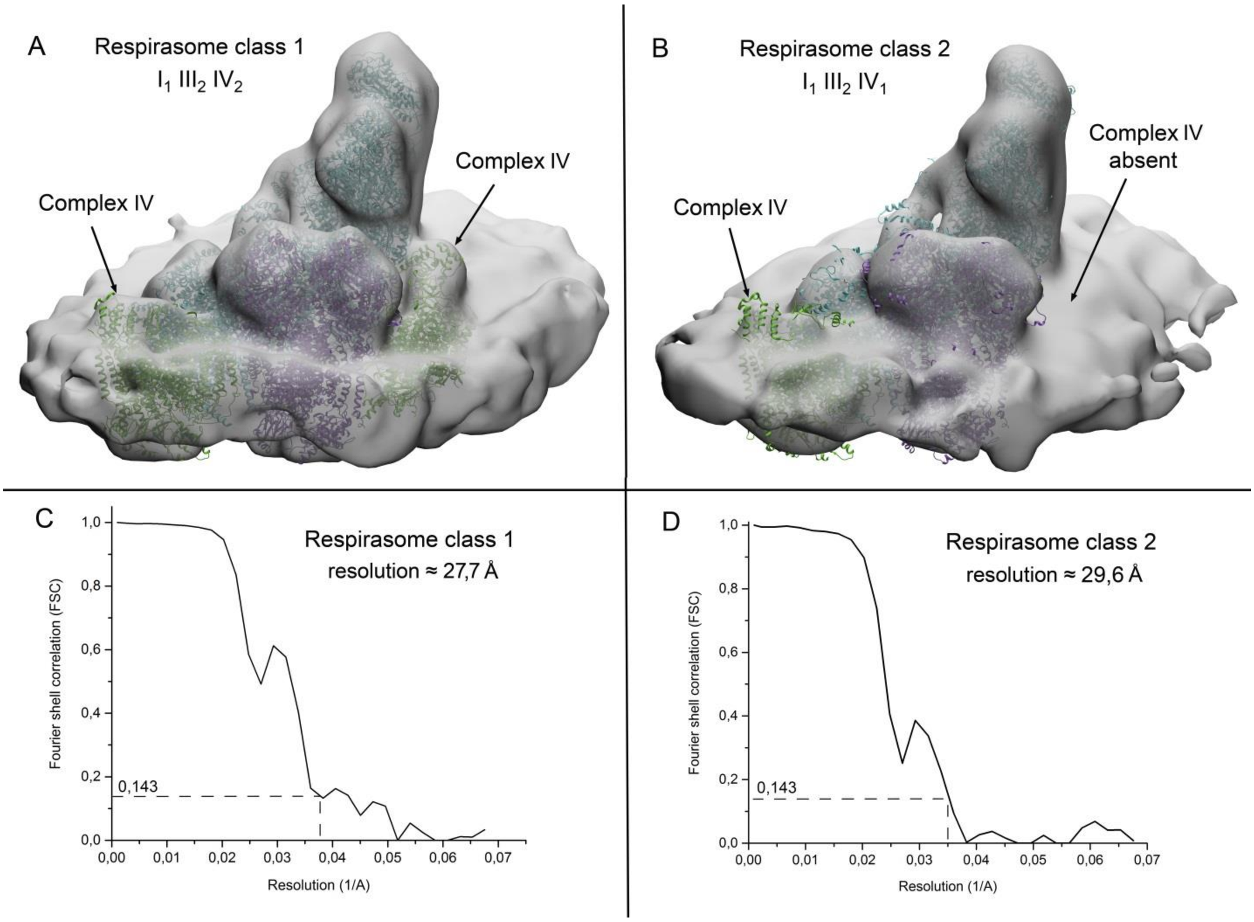


*Fig. S2. Two classes of respiratory chain supercomplexes. A. First class contains two complexes IV. B. Second class contains one complex IV. C-D. Fourier shell correlation graphs for resolution evaluation of class 1 and 2 correspondingly.*

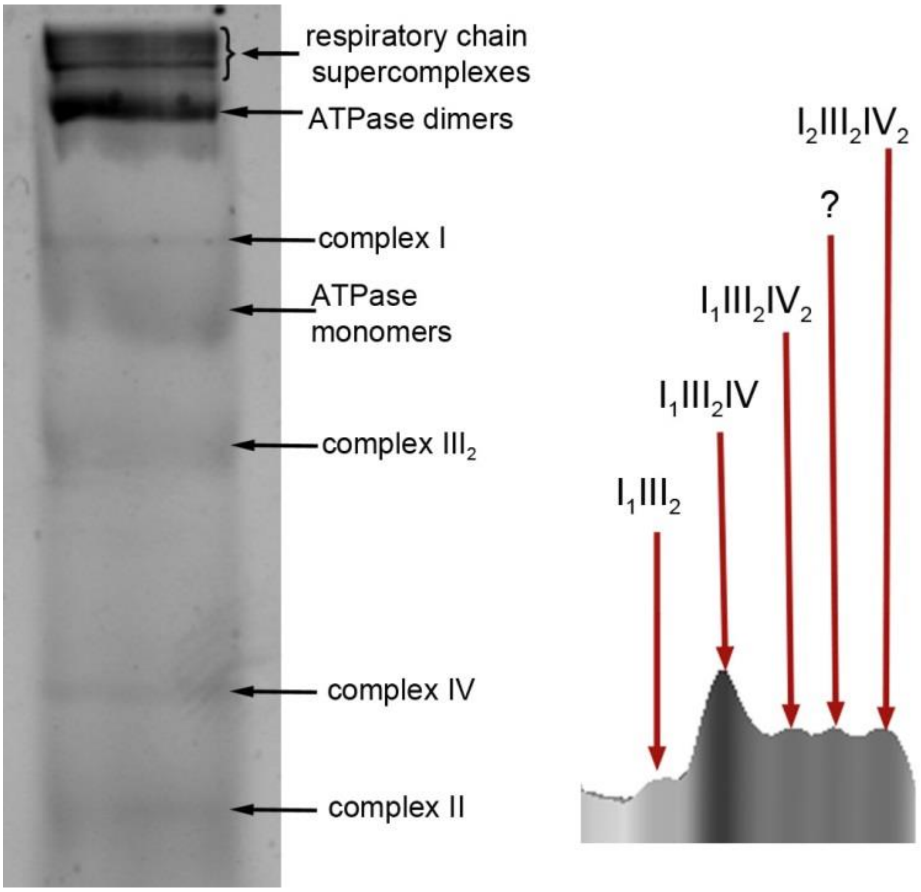


*Fig S3. Blue-native electrophoresis of mitochondrial fraction solubilized with digitonin and obtained from the same mitochondria that were used for cryo-EM experiments. On the right is a slice of a three-dimensional representation of a supercomplex-containing gel region. The height of the peaks in the 3D-reconstruction is proportional to the protein density in the gel (band brightness).*

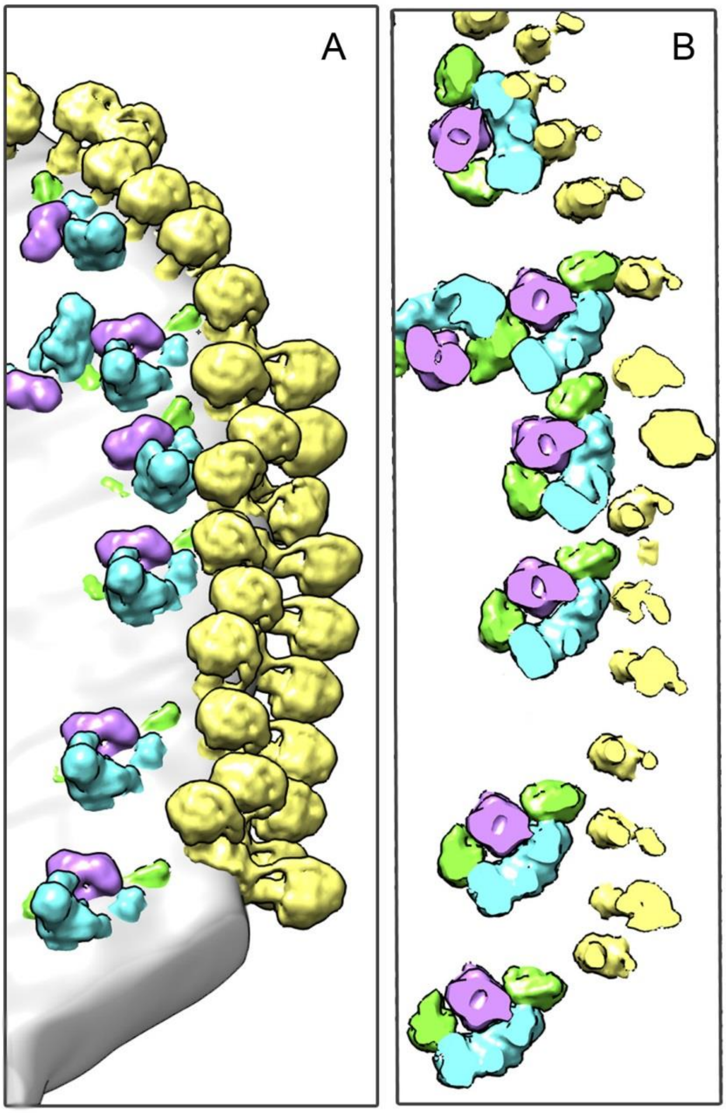


*Fig. S4. A. Surface rendering of a mitochondrial crista fragment containing linear hypercomplex of the respiratory chains and ATP synthases. B. Slice representing contacts of respirasomes and ATP synthases in the membrane. Colors are similar to Fig. 2-3.*

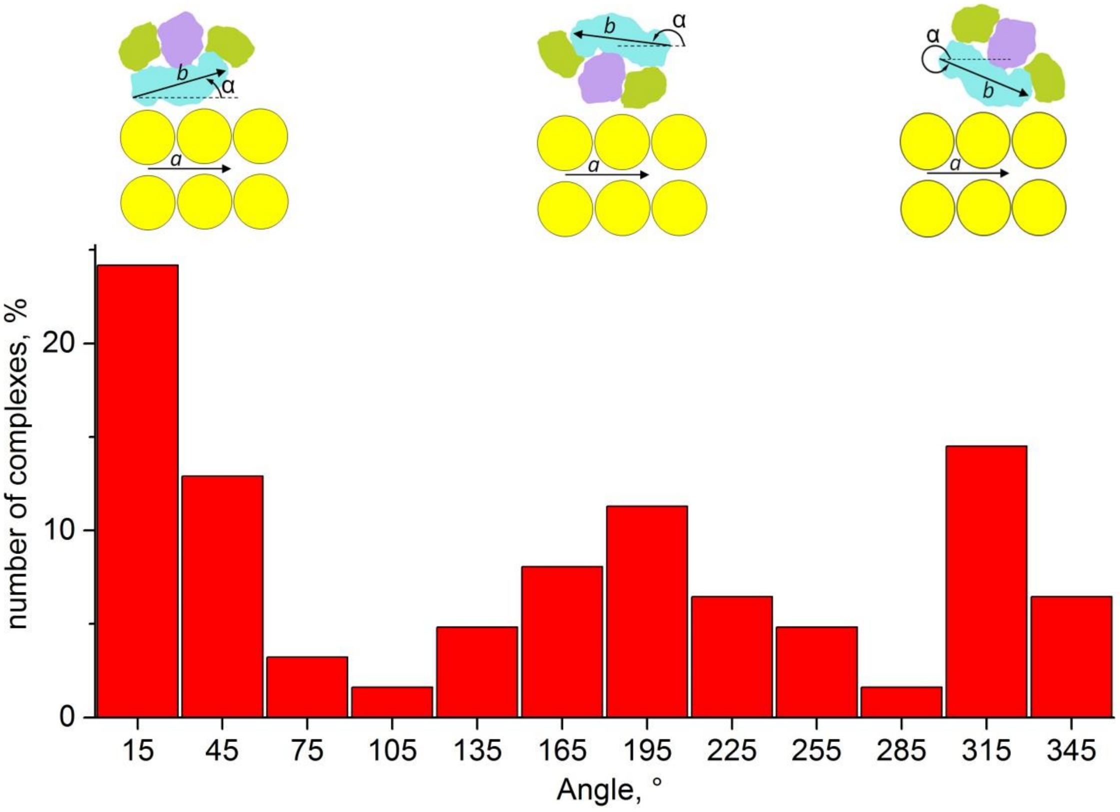


*Fig. S5. Statistical analysis of complexes I orientation relative to the rows of ATP synthases in the hypercomplexes. Only respirasomes from manually selected large hypercomplexes (like in fig.2, 3, S4) are included in this analysis. The inserts at the top show the approximate orientation of the respirasomes in the range of angles in the diagram below them. The direction of the ATPase row is determined from the orientation of the monomers in the ATPase dimer closest to the respirasome: vector* ***a*** *is defined as the vector product of unit vectors parallel to γ-subunits of ATPases. Vector* ***b*** *is directed along the membrane part of complex I (from ND1 to ND5 subunit along the longest interval). Angle α was determined as the angle between vector* ***b*** *and vector* ***a*** *projection to the membrane plane of respirasome. The final histogram is plotted with 30° step.*

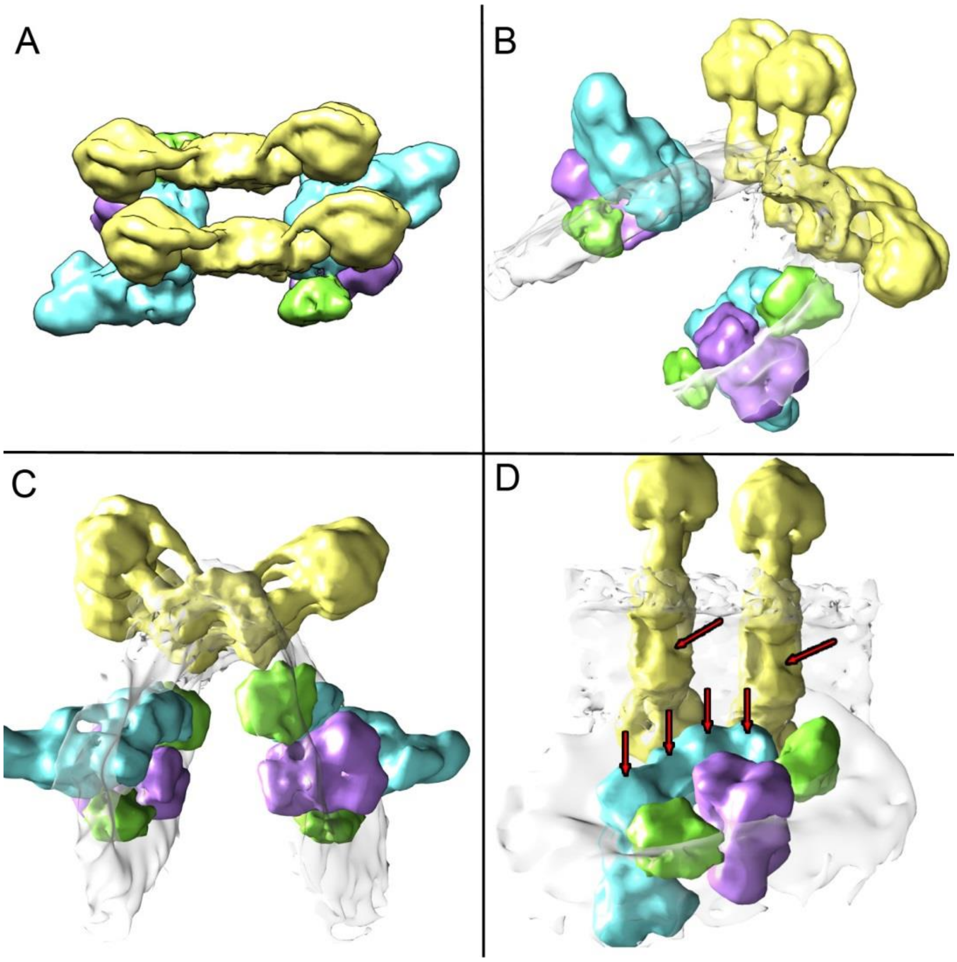


*Fig. S6. Reconstruction of mitochondrial crista fragment containing one symmetrical chain of the OXPHOS hypercomplex formed on the inner mitochondrial membrane fold. A-D. Various structure projections. In figure A the membrane is not shown, in B-D the membrane is shown transparent. In figure D only one respirasome is shown, arrows indicate the approximate location of proton pumps exits and ATP synthases' proton input channels.*

*Movie 1. Visualization of the placed-back crista reconstruction. Colors: yellow - ATP synthase, blue - complex I, purple - complex III dimer, green - complex IV.*
